# L-Regulon: A novel “soft-curation” approach supported by a semantic enriched reading for RegulonDB literature

**DOI:** 10.1101/2020.04.26.062745

**Authors:** Oscar Lithgow-Serrano, Socorro Gama-Castro, Cecilia Ishida-Gutiérrez, Julio Collado-Vides

## Abstract

Manual curation is a bottleneck in the processing of the vast amounts of knowledge present in the scientific literature in order to make such knowledge available in computational resources e.g., structured databases. Furthermore, the extraction of content is by necessity limited to the pre-defined concepts, features and relationships that conform to the model inherent in any knowledgebase. These pre-defined elements contrast with the rich knowledge that natural language is capable of conveying. Here we present a novel experiment of what we call “soft curation” supported by an ad-hoc tuned robust natural language processing development that quantifies semantic similarity across all sentences of a given corpus of literature. This underlying machine supports novel ways to navigate and read within individual papers as well as across papers of a corpus. As a first proof-of-principle experiment, we applied this approach to more than 100 collections of papers, selected from RegulonDB, that support knowledge of the regulation transcription initiation in *E. coli K-12*, resulting in L-Regulon (L for “linguistic”) version 1.0. Furthermore, we have initiated the mapping of RegulonDB curated promoters, promoters, to their evidence sentence in the given publication. We believe this is the first step in a novel approach for users and curators, in order to increase the accessibility of knowledge in ways yet to be discovered.

## Introduction

The pace with which scientific literature is being produced is clearly imposing new challenges to develop effective strategies that leverage the knowledge contained within thousands of publications. Content curation plays a crucial role in feeding pre-designed models of knowledge that support databases and, thus, in facilitating the finding of the more relevant objects [19]. It acts as a filter that separates the signal from the noise.

There is a growing number of curation strategies supported by Natural Language Processing (NLP) and Machine Learning (ML) tasks, and they have become a key source of information for bioinformatics repositories. Protein-protein interactions, regulatory interactions identification, entity association to ontologies, or even directed searches, are just some of aided curation examples [6, 17, 23]. It is important to emphasize that they are focused in facilitating access to specific information patterns. This kind of curation is vital and very helpful but it is designed on the premise that what is searched fits in a set of predefined and foreseen model and criteria, thus it only targets a fraction of the potential knowledge. This is the case of RegulonDB a database on transcriptional regulation in E. coli K-12.

RegulonDB ^1^ [7] is a manually curated standard resource, an organized and computable database, about the regulation of gene expression in Escherichia coli K-12. As a product of a continuous effort since 1991 it aims at integrating in a single repository all the scattered information about genetic regulation in this microorganism including elements about transcriptional regulation, such as: promoters, transcription units (TUs) and operons, transcription factors (TFs), effectors that affect TFs, active and inactive conformations of TFs, TFs binding sites (TFBSs), regulatory interactions (RIs) of TFs with their target genes/TUs, terminators, riboswitches, small RNAs and their target genes. It has also incorporated new concepts such as regulatory phrases of multiple TFBSs and GENSOR units (GUs). This information is limited to the main mechanistic elements of the classic model of transcription initiation and operon organization, with some extensions required to deal with the variants discovered through the years. These curated elements from each scientific publications (presumably roughly 20%), leaves out a vast amount of knowledge buried by hundreds of thousands of other statements and by the subtleness of natural language.

RegulonDB is the product of a tremendous maintained effort to organize the information conveyed in the curated articles. However, because natural language is highly complex, variable and amazingly expressive, to fit every single piece of this heterogeneous knowledge in pre-established structures is rather unfeasible. As a consequence, only a fraction of the information and knowledge contained in these articles has been curated in RegulonDB.

In order to alleviate this disconnection between the curated information –stored in RegulonDB– and the remaining unprocessed in the curated articles, we propose to link the curated objects in RegulonDB with their textual source by identifying the sentences that originated a curated piece of information and linking objects to sentences. Since these links locate the source sentences within their publications, this will directly provide the context for potentially broader analysis allowing a more comprehensive interpretation.

Furthermore, since scientific research is a social and cooperative product constructed iteratively and incrementally upon former knowledge, scientific articles are not self-contained pieces of knowledge, but new bits of information and knowledge that relies on and inter-connects with many others.

Despite the effort and good intentions of any author, due to the complexity of knowledge, of the enormous amount and the growing rate of published scientific literature in any field, it is improbable to consider all related research and access every single piece of relevant knowledge, particularly if its relation is not direct (e.g., a procedural or methodological relation but not on the same specific topic). Here is where we believe that computers and specifically automatic natural language processing can help to inter-connect similar conveyed ideas among a collection of articles.

We are here proposing a complementary semantic enrichment [3, 5], a “soft curation” focused in the whole publication textual content. It is based on, first, the association of already curated RegulonDB’s elements to their probable textual source within the publications, and, second, to interlink publications’ sentences by their semantic resemblance. The intention is that the resulting links feed a new browsing/reading strategy.

The goal of semantic enrichment is to improve utility, discovery and interoperability of content. It consists of enriching the original content with metadata, so that content could make more sense and to be better located in a broader context of related information for their consumers, including both humans and computers [3, 16].

There are several approaches to enhance the content’s semantics such as classification, tagging, categorization, indexing, etc. These strategies frequently include *Natural Language Processing* (NLP)tasks like Name Entity Recognition (NER), Syntactic Parsing, Word-Sense Disambiguation, Mapping with external Knowledge Organized Structures (KOS) (e.g. taxonomies, thesaurus, ontologies), Topic Clustering, etc.

But to the best of our understanding, none of the semantic enrichment strategies use semantic similarity in the way we do it: discovering semantically related sentences within a set of scientific articles and delivering those meaningful relations through links for each sentence. This functionality along with the links between RegulonDB’s objects and its source literature is what we call LRegulon (for Linguistic-Regulon).

We have developed this system in the interest of providing RegulonDB’s users new ways of interacting with information, to get a direct way to the source of the evidence of curated objects within their original context (source articles), and to track paths of research statements across publications, i.e., to seize better the time, effort and knowledge captured in scientific literature, curated or not.

We are still discovering the possibilities and sizing the aid that this kind of strategy can bring, but so far users have welcome very well the fresh perspective over the literature and knowledge.

## Materials and Methods

### 1 Methods

We had two main approaches: Inter-connect sentences in a set of articles through their meaning –by Semantic Textual Similarity (STS) [1, 8, 9]– and to link RegulonDB’s objects with their corresponding textual evidence within the literature. Each of these objectives were tackled with specific methods as described in the following.

#### 1.1 Interlink articles’ sentences through shared meaning

A possible approach to link sentences using their meaning could be linking phrases considered paraphrases among them. This would limit the universe of linked sentences to those having “exactly” the same meaning. Instead, we decided to discover suitable links measuring the semantic similarity of each sentence versus all others and keep the top N rated. On the one hand this brings more chances to find suitable sentences to link, and on the other hand it allows to discover related sentences that do not necessarily convey the “exact” same idea but an approximate one. Thus, it is the user who decides what to focus on: a closer meaning to the original sentence (higher score), or a more broader context similarity (a lower score).

We decided to apply a frequently used strategy consisting in applying several metrics which measure different similarity aspects of both sentences and then combine the scores into a single one [10, 21]. This strategy has proved to be robust to contexts’ changes. The strategy consists in the following steps:

1. Pre-process the text.
2. Apply multiple similarity metrics to each pair (STS step).
3. Combine the metrics in a single similarity score for each sentence.
4. For each sentence, select the top N related sentences.

The *preprocessing step* consists in identifying multiword object names by obtaining a list of multiword objects names from RegulonDB, sort them (from longest to shortest), and look for occurrences within texts by using the left-right longest match; and next to lemmatize lexical units, tag Part-of-Speech (POS) and obtain the dependency trees^2^.

Three types of similarity metrics were included in the *STS step*: string metrics, which compare how similar are the words and their order between two sentences; distributional metrics, which are based on the premise that context is representative of word meaning, so words are represented as multidimensional vectors into geometric spaces where proximity is used as the SS; and ontological metrics, which rely on the knowledge’s structure to determine how close is the meaning of two concepts.

A total of eight metrics were used:

1. Ontology-based measure, using the common vector technique, Wordnet as the knowledge source, and Lin98 metric.
2. Normalized Levenshtein.
3. Jaccard.
4. N-Gram.
5. Averaged word embedding, using pretrained GLOVE vectors.
6. Averaged word embedding, using ad hoc vectors trained over literature of transcriptional-regulation.
7. Sentence embedding from common vector using ad hoc vectors trained over Transcriptional-Regulation literature.
8. Ontology-based measure, using domain-dependent ontology and Lin98 metric.

The final *combination step* consists of using regression and ensemble algorithms to merge the individual results from the previous step. The regression algorithms to be used are Linear Regression, Multilayer-Perceptron, and Random Forest, whereas the ensemble algorithms are Bagging and Voting.

A complete description of each of the individual metrics, regression and ensemble algorithms can be found in [12].

#### 1.2 Other approaches to leverage Semantic Similarity

Besides using the *semantic similarity* (SS) among the sentences to generate the interlinks we implemented two other novel approaches to leverage the already computed SS. Below their aims and methods are described.

##### Semantic network of publications

In the interlinking process, sentences are linked not only to other sentences within the same publication but also to sentences in other publications of the collection (external links). We used this external links to build a semantic directed graph where the nodes represent the publications and the edges the SS strength among publications. Because it is a directed graph, the edge strength between publications *A* and *B* is computed adding the similarity scores of all the interlinks that connect sentences in A with sentences in B, and the *B* -*A* edge comes from the interlinks that connect sentences in B with sentences in A.

This graphs provide a comprehensive overview of one collection of publications. It aims at providing an analysis tool so the user, through a quick view, can detect which publications are more related to all the others, which ones are more isolated and which ones have the strongest semantic connections. It is worth noticing that this meta-analysis should be interpreted accordingly to the collection. For example, if the collection is constituted of a specific topic, then, review articles –even if not officially classified as reviews– should be identified as the more interconnected; and because the graph is directed, publications dependencies –even if not cited– can also be detected.

##### Extractive summary based in Semantic Similarity

Another novel exploit of SS is to use it as the foundation of an automatic publication summarization process. For this task only the interlinks within the same publication are used. The aim is to score the centrality (eq. 1) of each sentence within the article based in its semantic relation with the other sentences. The score is used to sort the sentences in descending order and, finally, the top N sentences are selected accordingly to the desired summarization percentage.

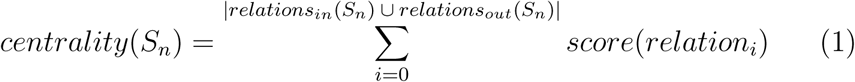

This approach is inspired in the Textual-Energy summarization technique [2, 4] which uses common words between a pair of sentences (*A* and *B*) as well as the common words between *A* and *C* and between *C* and *B* to compute the weight of a sentence in the document. This is in turn a derivation of the centroid-based summarization technique [18].

A differentiating feature is that our proposal is not based in textual co-occurrences but on semantic ones, i.e., the sentences which its meaning is more repeated through the document are very probably those that condensate most of the key information.

This summarization process produces extractive^3^ summaries, that are formed by selecting and putting together a subset of sentences that cover as much information as possible from the publication.

#### 1.3 Link RegulonDB objects with supporting textual evidence in literature

The aim here is to do a reverse interlinking, that is, starting from the RegulonDB objects, try to identify those sentences within the publications that could be the curation source of the object and its properties.

Through years of experience, curators have noticed that particular patterns occur in the sentences that state the information that is curated in RegulonDB’s objects. Leveraging curators experience and knowledge, we collected a set of heuristic rules based in specific words and syntactic patterns. Basically the strategy is to identify those sentences that match one or more rules and to assign them a score based in the number and co-appearance of identified object properties within the sentence. These rules rely on words and patterns that are specific to the type of object for which we are trying to find its curation sources, thus a different set of rules have to be manually crafted for each RegulonDB object type. Therefore we approached this task by stages, one object type at a time.

So far we implemented the reverse interlinking only of promoters. The properties of promoters curated in RegulonDB are: absolute+1 position, -10-box, -35-box, sigma factor and evidence of the transcription-initiation start site.

Next we describe the rules defined for each of these properties.

##### Absolute+1 position

To match sentences that express this property they have to full fill one of the following conditions:

- Contain the term *5’* and, *mapping* or *determination*.
- Contain *TSS* and optionally *promoter*, but do not contain the word *solution*.
- Contain *transcriptional start site* and optionally *determined*, and prefer those that do not contain any of TFs names.

The previous rules are mandatory in order to tag the sentence as *absolute*+*1 position*. Additional rules that can increase the confidence tagging score based in the coappearance of other properties in the same sentence are:

- If the sentences contains *absolute*+*1* position with the *5’* term and information about *evidence of transcription-initiation*
- if the sentences contains contains *absolute*+*1* position with the *TSS* term and information about *evidence of transcription-initiation* specifically mentioning *primer extension* or *S1*

##### -10 box

Mandatory one of the following conditions:

- The term *-10* and one of the following terms: *hexamer, Pribnow, box, promoter, transcription, consensus, region, sequence, site, element*; but do not contain any of the terms: *Primer, ribosome, ribosomal, discrimination, shine, operator, spacer, -100, -1000, repeat, NarL, NanR*.
- Contains any sequence of bases (AGTC) with a size between 6 to 8 and any of the terms: *-10, -35, hexamer, Pribnow*; but do not contain any of the terms: *Primer, ribosome, ribosomal, discrimination, shine, operator, spacer, -100, -1000, repeat, NarL, NanR*.

Optional coappearances with other properties that increase the confidence score:

- The *-10 box* along with the *-35 box* and the *sigma factor*.
- Or the *-10 box* along with the *-35 box* and the *evidence of transcription-initiation*.
- Or the *-10 box* along with the *-35 box* specifically mentioning a base sequence of size between 6 and 8 pb.
- Or the *-10 box* along with the *-35 box*.

##### -35 box

Mandatory one of the following conditions:

- The term *-10* and one of the following terms: *hexamer, Pribnow, box, promoter, transcription, consensus, region, sequence, site, element*; but do not contain any of the terms: *Primer, ribosome, ribosomal, discrimination, shine, operator, spacer, -100, -1000, repeat, NarL, NanR*.
- Contains any sequence of bases (AGTC) with a size between 6 to 8 and any of the terms: *-10, -35, hexamer, Pribnow*; but do not contain any of the terms: *Primer, ribosome, ribosomal, discrimination, shine, opera, spacer, -100, -1000, repeat, NarL, NanR*.

Optional coappearances with other properties that increase the confidence score:

- The *-35 box* along with the *-10 box* and the *sigma factor*.
- Or the *-35 box* along with the *-10 box* and the *evidence of transcription-initiation*.
- Or the *-35 box* along with the *-10 box* specifically mentioning a base sequence of size between 6 and 8.
- Or the *-35 box* along with the *-10 box*.

##### Sigma factor

Mandatory one of the following conditions:

- The term *sigma*, one of the following patters *consensus, dependent, promoter, transcript*\*, *by* [*the associated sigma*], and the gene associated to the promoter we are dealing with.
- Contains any of the following terms *s70, s32, s28, s19, s38, s54, s24, RpoS, RpoN, RpoD, RpoE, FecI, FliA,RpoH*.

##### Evidence of the transcription-initiation (TrInMa)

Mandatory one of the following conditions:

- Mention the phrase *primer extension* and optionally any of the verbs *map, determine* or *perform*.
- Contains the phrase *race* and optionally any of the verbs *map, determine* or *perform*.
- Contains the term *S1*, and one of the patterns *nuclease, S1 protection, S1 analysis*, or one of the verbs *map, determine* or *perform*; but do not contain the phrases *table S1* or *figure S1*.

## Results

In this section we first present the results of the two main enrichment approaches and then the results of the system as a whole.

### 1.4 Interlink articles’ sentences through shared meaning

To the best of our knowledge, this to read/navigate through collections of literature has not been described before.

In the following subsections, first the quality of the semantic links is tackled and next a brief summary of the interlinked collections is presented.

#### 1.4.1 Quality of the semantic links

The quality of the semantic-interlinks was evaluated indirectly by measuring the precision of the similarity metrics. This was done through a 10-fold cross-validation over a Similarity Corpus [13], which is a graded textual similarity corpus that was specifically designed to be used for training similarity models.

The corpus consists of 171 pairs of sentences extracted from transcriptional regulation literature (5, 600 articles). Each pair of sentences was rated with an ordinal scale ranging from 0 to 4 – where 4 means that both sentences express the same meaning and 0 that their meanings are completely different– by 3 annotators taken by chance from a group of 7 (non-fully-crossed design). The resulting corpus has a *very good interagreement* [25] coefficient (Gwet’s AC2) of 0.8696.

Table 1 summarizes individual metrics scores, and regression and ensemble performance is presented in table 2. From the data the best individual metric with a *ρ* of 0.604 is the one using GloVe embeddings trained on transcriptional-regulation literature, whereas the best merge strategies were Random Forest and Voting with a *ρ* of 0.683 and 0.693 respectively. These correlation scores are considered high [15] given how specialized is the literature vocabulary and how complex are some of its sentences compared to general domain literature (news, web, etc.). Further details about the implementation, evaluation and analysis can be found in [12].

**Table 1.** Correlation by measure

| Measure | Correlation |  |  |
| --- | --- | --- | --- |
| | $\rho$ | $r_s$ | diff |
| 1) Wordnet | ○ 0.261 | 0.277 | 0.016 |
| 2) Levenshtein | 0.236 | 0.269 | 0.033 |
| 3) Jaccard | 0.484 | 0.524 | 0.039 |
| 4) NGRAM | 0.241 | 0.257 | 0.015 |
| 5) Glove (averaged) | 0.391 | 0.435 | 0.044 |
| 6) AdHoc-Glove (averaged) | 0.424 | 0.540 | 0.116 |
| 7) AdHoc-Glove (common-vector) | ● 0.604 | 0.634 | 0.030 |
| 8) AdHoc-ontology | 0.484 | 0.484 | 0.000 |

**Table 2.** Regression and ensembles performance

| Algorithm | Correlation | St. dev. |
| --- | --- | --- |
| Linear Regression | 0.622 | 0.15 |
| Random Forest | ● 0.683 | 0.14 |
| Perceptron | 0.642 | 0.18 |
| Bagging | 0.691 | 0.14 |
| Voting | ● 0.693 | 0.13 |

#### 1.4.2 Interlinked collections

The interlinking process is very intensive because it evaluates the similarity between each sentence and all the other sentences from all other publications, i.e. it grows exponentially in relation to the number of sentences/publications. For this reason it is suggested to be applied on collections of no more than hundreds of articles. Because there is no restriction for a publication to be included in more than one collection, as many collections as needed can be generated to cover different analysis purposes. Collections can be seen as viewpoints over the literature; for example, a collection of promoter publications can be interlinked for an element type analysis, whereas an analysis around a specific transcriptional regulator would use a collection formed by articles related, for instance, to OxyR.

So far we have processed and interlinked the collections reported in the table 3, all containing articles obtained from RegulonDB literature. Another 111 collections, each one corresponding to a *Gensor Unit* (GU), have been also processed. Literature for each GU groups all papers cited in RegulonDB for all objects regulated by a given transcription factor. Together, the GU collections have 559 publications, 166, 425 sentences and 821, 000 relations. Version 1.0 of L-Regulon can be accessed at http://132.248.34.183:81.

**Table 3.** Interlinked sets.

| Collection | Articles | Sentences | Relations |
| --- | --- | --- | --- |
| Ris rob | 36 | 13,131 | 96,396 |
| Ris paax | 17 | 5,610 | 20,781 |
| Regulon NarP | 73 | 18,916 | 236,349 |
| Regulon SoxR | 64 | 14,674 | 179,279 |
| Regulon AraC | 54 | 11,955 | 130,893 |
| Anaerobiosis ModE | 6 | 1,209 | 97,195 |
| Anaerobiosis FNR | 10 | 1,943 | 16,1917 |

### 1.5 Other approaches to leverage Semantic Similarity

#### 1.5.1 Interactive semantic network of publications

Figure 1 shows the network of the collection *Regulon AraC*. The nodes are labeled with the publication’s PMID and, in this example, the graph is organized in a concentric layout but there are other 7 possible layouts to fit the user preferences. There are other graphical hints to facilitate the analysis: the edges width and color are related to the strength of the semantic relation; the node color is related to the number of interlinks it receives (the darker the color the more interlinks it receives); and, when a node is selected its interlinks and connected nodes are highlighted whereas all others are toned down.

**Figure 1.**
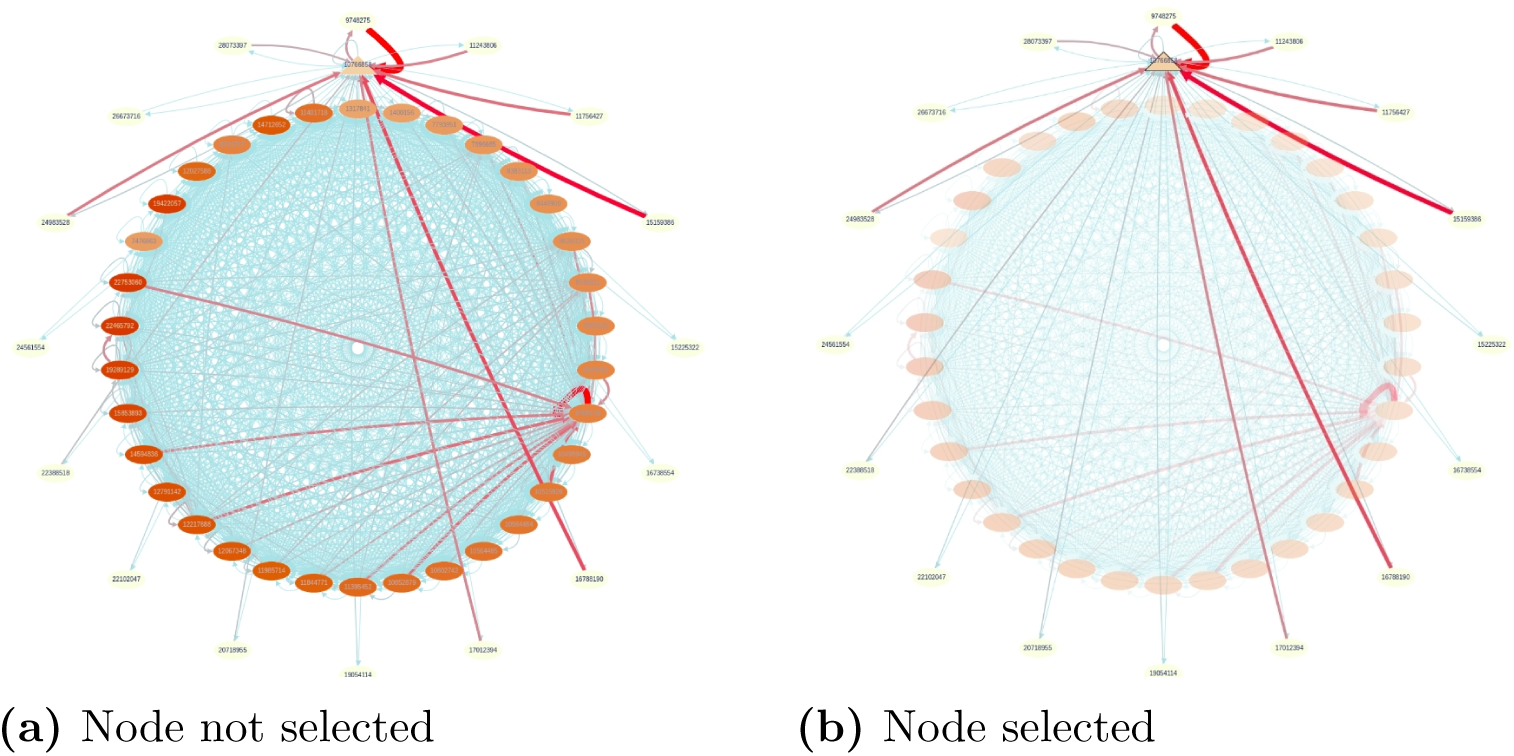
Semantic network of publications of *Regulon AraC* collection.

#### 1.5.2 Extractive summary based on the sentences’ similarity centrality

As shown in fig. 2 when summarization is activated the selected sentences are highlighted both in the publication text and in the eagle-view where it can be appreciated the section of the article where the sentences are coming from.

**Figure 2.**
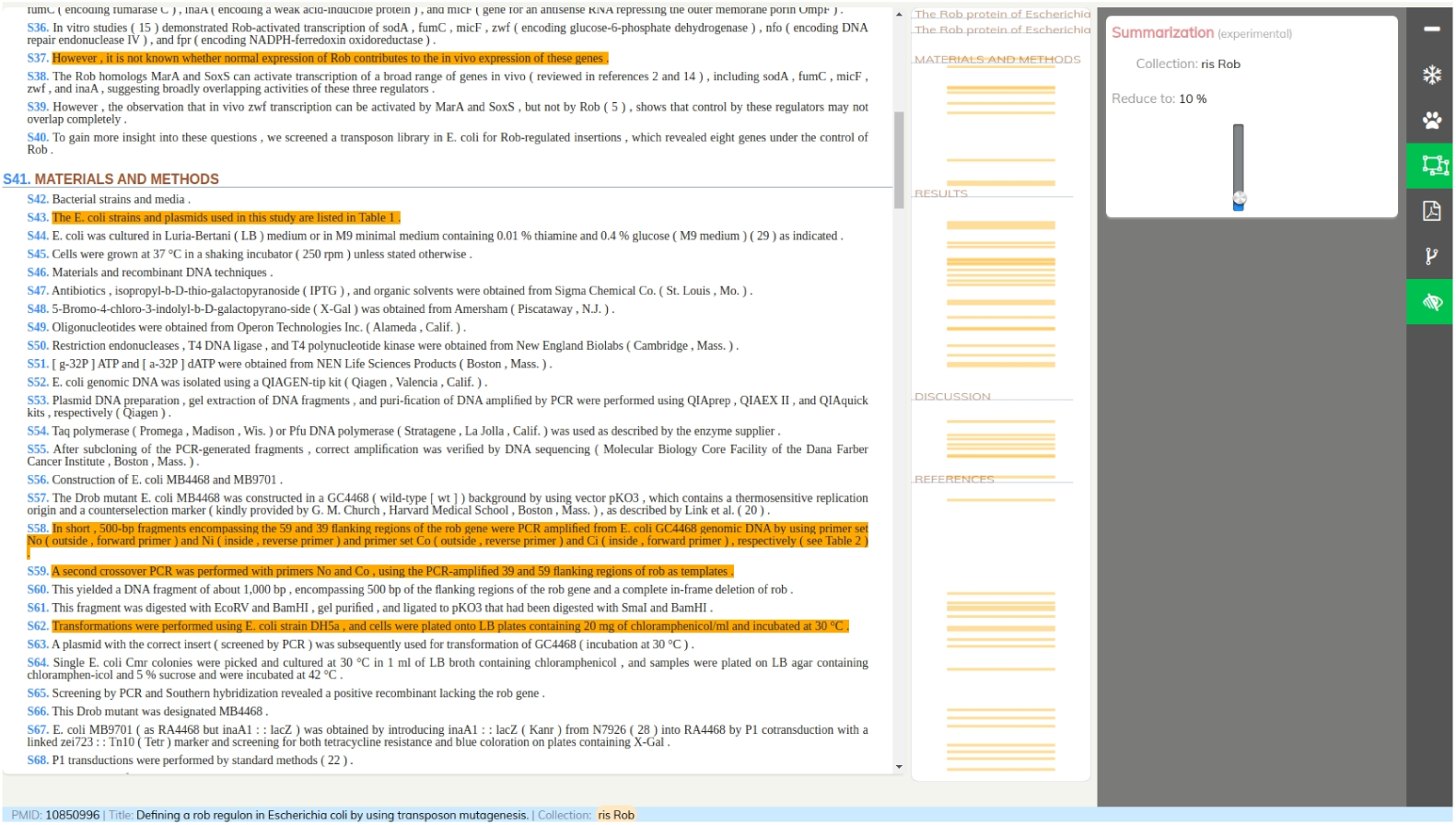
Extractive summary consisting of 10% of the publication’s sentences.

### 1.6 Linking RegulonDB objects with supporting textual evidence in literature

In order to provide flexibility and prepare the system for the next stages (other object types) a meta-syntax was designed to express the rules distilled by the curators. The syntax is based in the YAML syntax and a yaml rules file have to be created for each object type. Thus, for the promoters, a *promoter*.*yaml* file must contains the rules for each of the properties of the Promoters objects.

The next box has an example of the file structure where the *absolute*+*1 position* property is defined. It consists of a set of *patterns* and a set of *post scoring* conditions.

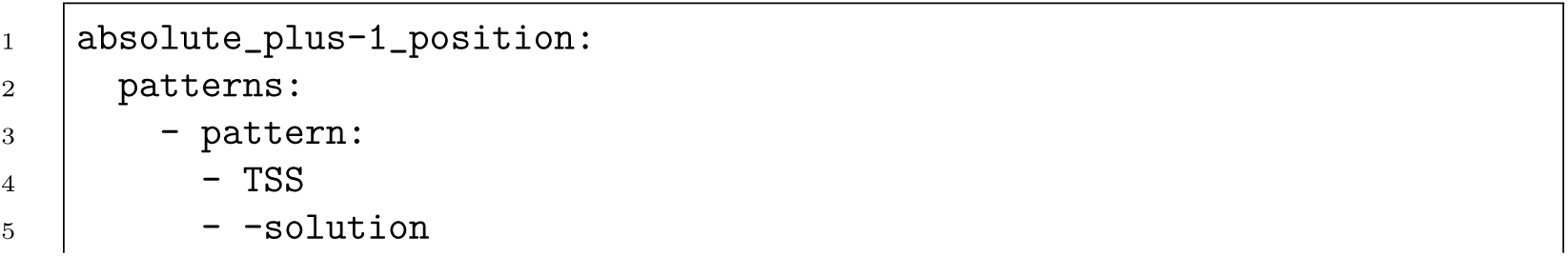
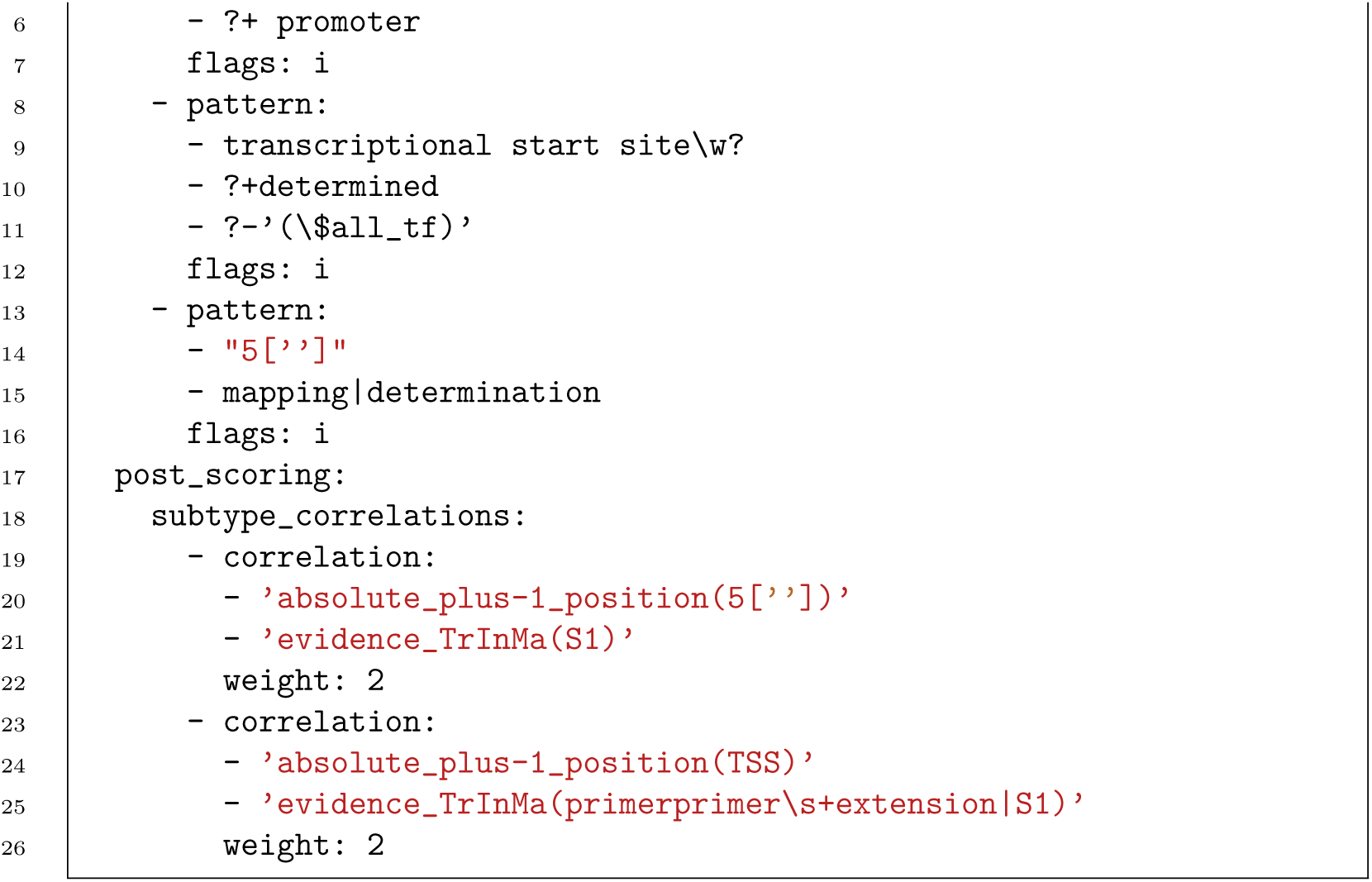

The patterns section can have one or more mandatory *pattern*s each of which is formed by one or more conditions. A condition can be a word (e.g. TSS), a regular expression (e.g. transcriptional start site \w?) or a metavariable (e.g. $all tf). At the runtime the metavariables are substituted by regular expressions that represent list of values extracted from RegulonDB. Currently, the available variables are *referred_promoter, referred_sigma* and *referred_gene*, which respectively represent the list of all TF names, promoter names, sigma names and gene names associated with the publication being processed. Conditions are considered as required by default but if preceded by ‘?’ then they become optional. An optional condition can be qualified as positive (i.e. preceded by ‘?+’) or negative (i.e. preceded by ‘?-’), positive conditions increase the confidence score while negative conditions decrease it. Finally, if a condition is not optional and is preceded by ‘-’ that indicates that the sentence must not contains the condition.

The *post scoring* section holds the correlation rules among the property being defined, in this case the absolute+1 position, and the other properties. These rules are used to increase the confidence score when certain properties appear together in a sentence. In the example the first correlation rule expresses that if the property *absolute*+*1 position* with the term *5’* is found in a sentence that also has the *evidence-TrInMa* with the term *S1* then the confidence score is multiplied by two.

#### Applying the rules

To avoid ambiguity we focused in those publications that refer to only one promoter. To obtain those publications we first got the list of promoters from RegulonDB along with their associated PMIDs. Then, those PMIDs linked to more than one promoter were filtered out. Next, we obtained the PDFs of the remaining PMIDs and extracted the textual content sentences with our PDFTxtextractor tool which has the capacity to extract the section in the paper (abstract, introduction, methods, etc.) where the sentence occurs.

In order to maximize the probability of finding a curation source sentence in the original articles and not only references on posterior publications, the rules were applied to sentences that do not belong to the sections abstract, introduction or references. This could be done thanks to our pdf-extraction tool which in the majority of the cases can identify the section of the article where the sentence occurs.

Finally, the set of regular expression rules was applied to each of the selected sentences. Based on the number of fulfilled patterns a confidence score was computed for each assigned tag in each sentence. Only those sentences with non-zero score were reported as source curation sentences for the promoter associated to the publication being processed.

This task is done as an off-line process that stores the found reverse links within the system’s database, thus when requested they are only retrieved from the database.

#### Found data

As a result of applying the rules we identified 5, 426 source sentences within 587 articles. The sentences expressed properties of 508 different promoters with the following distribution: 2, 405 sentences had evidences of *TrInMa*, 1, 326 expressed *sigma factors*, 1, 016 and 1, 008 sentences mentioned *-10 box* and *-35 box* data respectively, and 516 sentences stated the *absolute*+*1 position*. About properties co-appearance, 811 sentences expressed more than one property, 32 contained three properties and only 2 sentences were found with four properties. The vast majority of properties co-occurrence was *-10 box* with *-35 box* with 660 cases followed by 80 cases of *absolute*+*1 position* with *evidence TrInMa*.

#### The system interface

The ultimate goal of this feature is to help the user to find the most probable original-sentences from where the RegulonDB object was created. The discovered sentences associated to an object are displayed as shown in fig. 3. It is organized in a top-down layout; first the publication title, authors and journal is shown, below the list of all the object properties found in that publication and, nested in each property, the list of sentences that fulfilled the property’s rules.

**Figure 3.**
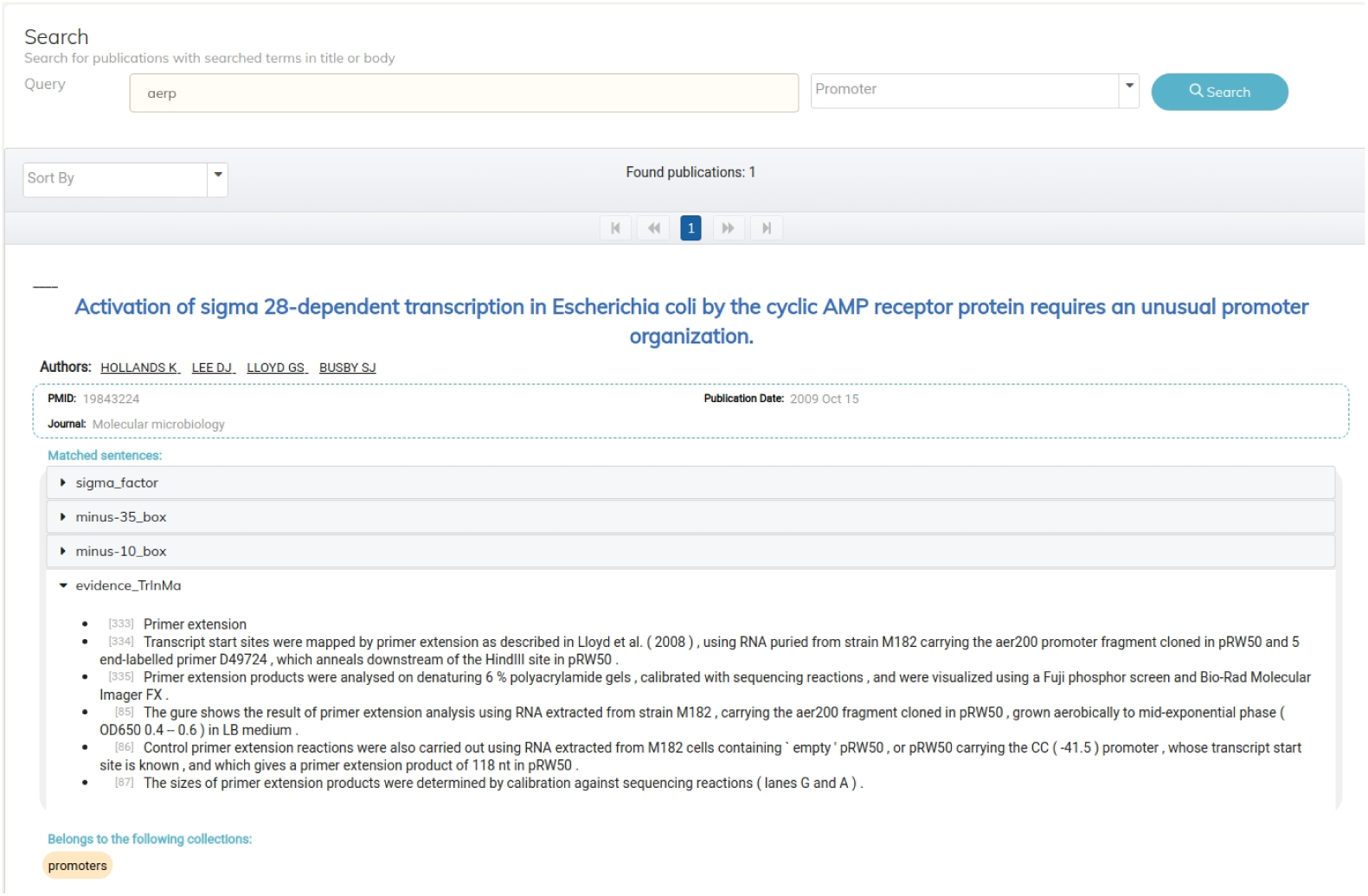
GUI of reverse link feature.

This information can be accessed through two entry points: by a direct search within L-Regulon where user must specify the name and type of the searched object (fig. 3), or by a link activated from RegulonDB object page that would open the L-Regulon screen.

### 1.7 System results

In order to grasp the kind of help that LRegulon can bring to RegulonDB users, here we present three examples (tables 4, 5, 6) of sentence interlinking along with a transcriptional-regulation expert interpretation.

**Table 4.**
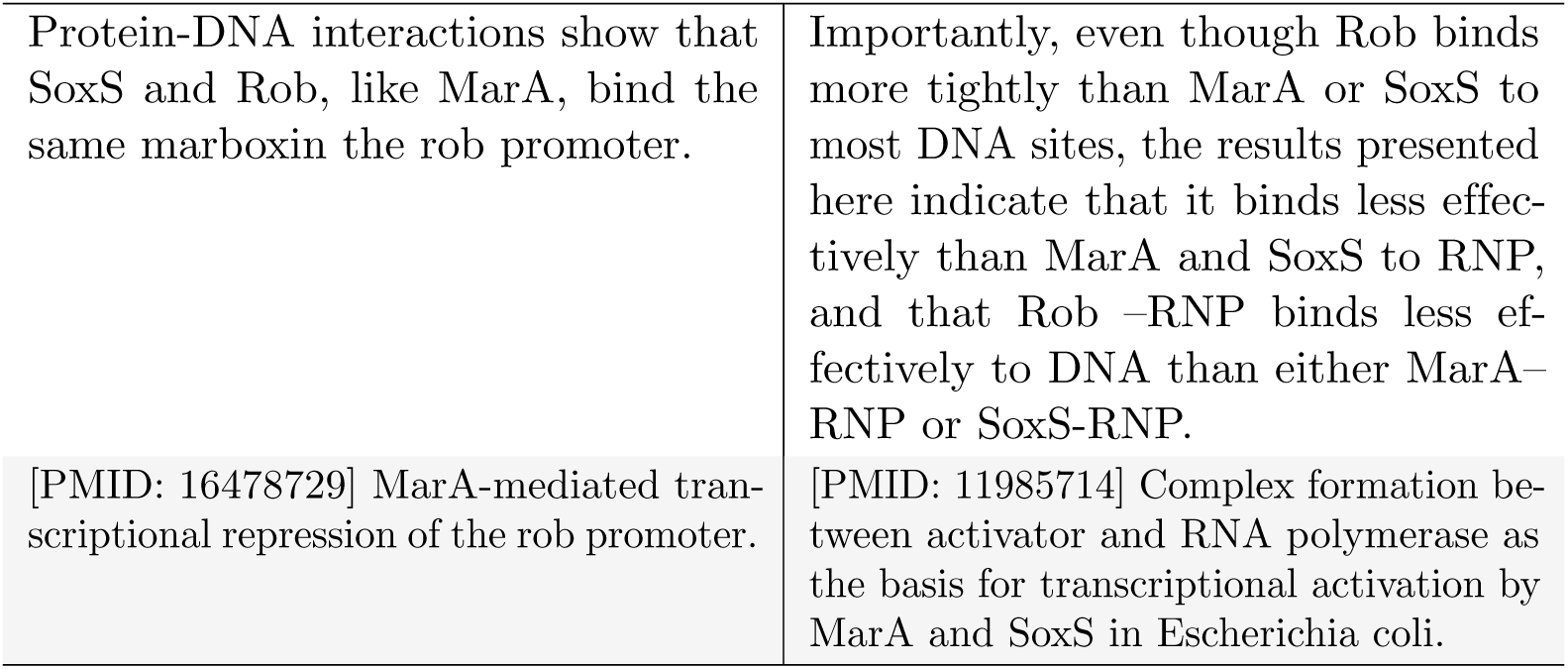
Example 1. Interlink example 1, the first row contains the sentences and the second the article title; left column is the origin publication and the right one the target.

**Table 5.** Example 2. Interlink example 2, the first row contains the sentences and the second the article title; left column is the origin publication and the right one the target.

|  |  |
| --- | --- |
| Protein-DNA interactions show that SoxS and Rob, like MarA, bind the same marboxin the rob promoter. | Likewise, MarA, SoxS, and Rob are all repressors of the soxS and rob promoters. |
| [PMID: 16478729] MarA-mediated transcriptional repression of the rob promoter. | [PMID: 22753060] Transcriptional cross talk within the mar-sox-rob regulon in Escherichia coli is limited to the rob and marRAB operons. |

**Table 6.**
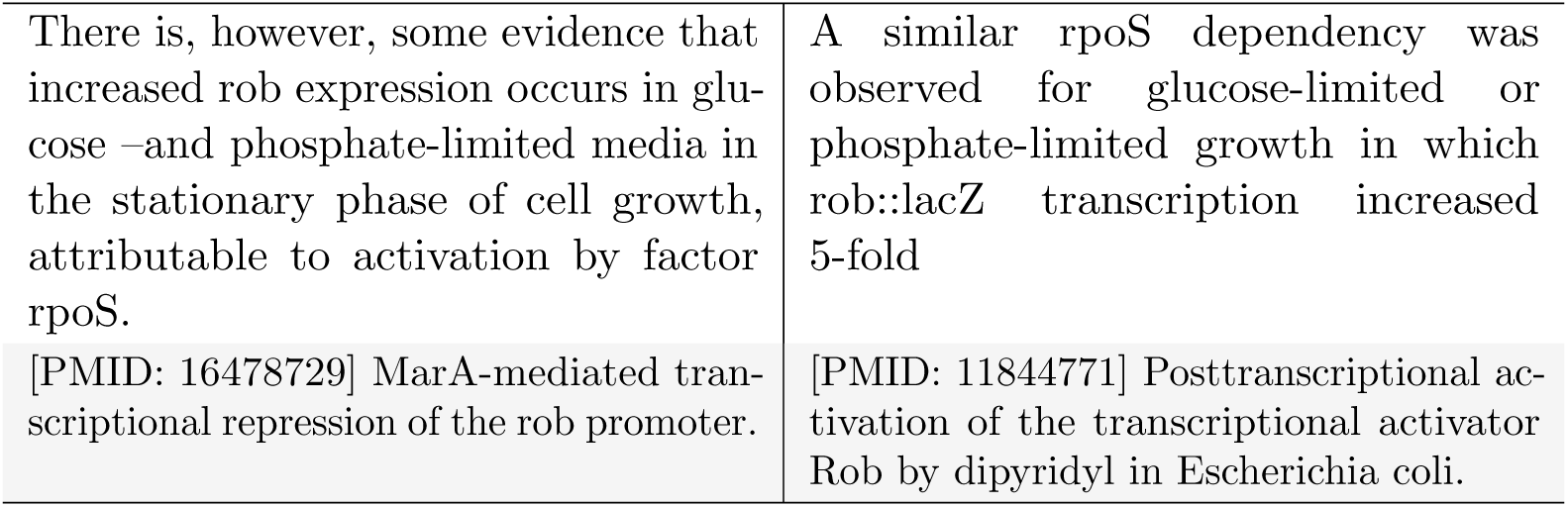
Example 3. Interlink example 3, the first row contains the sentences and the second the article title; left column is the origin publication and the right one the target.

In summary version 1.0 of L-Regulon here presented contains 118 interlinked collections, with 111 related to GENSOR Units plus 7 more. In detail that means more than 800 articles, 233k sentences and almost 1.8 million of relationships (interlinks). This data also feeds the semantic network and extractive summary features. This version also contains links between 508 promoters in RegulonDB and their source sentences.

## 2 Discussion

With the aim of enriching the textual content of RegulonDB’s publications, we have proposed and implemented a new soft-curation approach which produces a network of semantically related sentences, and a browsing/reading strategy derived from it. These assets are made available through a set of tools (interlink browsing, automatic summarization, and interconnected publications graph) disposed in a web-based application that completes the content curation pipeline by implementing the sharing element over both, the automatic produced semantic associations and the manually curated associations done by experts in the Lisen-&-Curate system.

Semantically enriched content, also known as smart content, lets you find topically related content and then recombine it to create new perspectives over data for specialized audiences, knowledge environments or learning objectives. It can also be considered a content curation strategy of the mashup type. Mashups are curated juxtapositions where merging existing content is used to contextualize and create a new point of view over the information.

This kind of curation does not impose a specific and foreseen targeted goal. Instead, it offers a stepping stone for other curation processes, either from human or computational processes. Another key idea is that within content curation seeking and making sense of the curated information represents only 2/3’s of the task, the third element is sharing; this is what allows others to access the added value produced over the enriched content. Particular attention is paid to this through its innovative web-based application where users can browse RegulonDB’s publications and exploit the curated interlink-semantic-network by different means and, also, have access to the source sentences of RegulonDB’s objects. This last feature helps to close the gap of RegulonDB and the FAIR principles because it makes the source article sentences readily available, promoting scientific transparency and accessibility.

It is worth noticing that LRegulon features are not dispersed tools but NLP-powered steps within a literature exploratory reading pipeline. To show this integration below we describe a possible LRegulon’s features use-sequence and their motivation.

First, users can look over the *interlinked network* at the level of the publication. This provides graphical insights of how a cluster (a content viewpoint) is interconnected and how some publications are more related to others by their semantic similarity. Then, the user can select that publication which receives or from which more connections depart and go directly to the LRegulon *content reader module*. There, a processed version of the paper next to its PDF is available for reading. During the reading, the user has access to a *summarization tool* that highlights a defined percentage of the most “central” sentences, determined from the number and strength of semantic links to other sentences within the same article. Finally, while skimming the summary (highlighted sentences), the user can find a statement relevant to his purposes and activate the *interlinking tool* to access other sentences with similar meanings. These interlinks make it possible to quickly find related content and perform a transversal reading (across articles) guided by semantics.

In this research, we have faced fundamental questions about the semantical enrichment environment that fits the needs of RegulonDB-literature users. Some of the inquiries are: What types of enrichment can be helpful to curators and general content consumers of RegulonDB? What are the benefits and returns of investment in developing these environments? How are we delivering the enriched content to final users? Which external building knowledge blocks or resources (i.e., thesaurus, taxonomies, ontologies, corpus, etc.) do we need to process the content? Some of these are still open questions that will be only be answered after a considerable period of utilization of the system.

## 3 Conclusions

The aim of this paper is to propose a novel literature exploratory reading tool that improved the utility, discovery and interoperability of collections of related scientific articles. We have tackled this challenge by an automatic mash-up curation strategy based on the identification of the object’s curation-source-sentences and the semantic interlinking of publication statements. We have also proposed to use the sentence-semantic-similarities to generate a directed graph of semantically connected publications and an extractive summarization strategy. To the best of our understanding, no other semantic enrichment strategy uses semantic similarity in this way.

This research was concerned with RegulonDB literature and audience; however, the system is certainly applicable to other research domains, and in general to any collection of related documents.

Thus far the analysis does not enable us to determine the long-term value of this new reading approach. However, we believe that will be a starting point for new research lines on the subject of NLP-assisted knowledge-exploitation.

Future work would focus on performance improvements and new enrichment strategies. As to the performance, enhancements will involve accelerating the STS computation to be able to interlink larger collections. An attractive alternative for this is the use of pre-computed sentence embeddings [11, 20, 22, 24].

Regarding new ideas, it may involve two approaches: connect users and content and, integrate other enrichment strategies. *Connect users and content* could include the creation of user profiles to hold user preferences, discovery pathways (starred sequence of traversed interlinks), topics of interest, and automatic user-collaboration-suggestions based on similarities of individual profiles (topics of interest, discovery pathways, etc.). New types of enrichment could include a *chronology curation* which would consist of connecting sentences across papers but sorted by the publications’ dates and, *distillation* which would consist of automatically-generated simplified texts.

## Acknowledgments

We acknowledge useful discussions with several members of the laboratorio, and acknowledge computational support by Víctor Del Moral and César Bonavides. We acknowledge funding by UNAM, and by NIH grant number 5RO1-GM110597-04 and 1RO1GM131643-01A1. J.C.-V. acknowledges support by DGAPA from UNAM during his sabbatical leave at the Center for Genomic Regulation in Barcelona.

## Footnotes

1 http://regulondb.ccg.unam.mx/

2 We used the Stanford CoreNLP toolkit [14]

3 In this type of summarization sentences are not reformulated nor syntactically adapted.

